# Prelimbic cortex neural encoding dynamically tracks expected outcome value

**DOI:** 10.1101/2022.05.18.492483

**Authors:** Mark Niedringhaus, Elizabeth A. West

## Abstract

Animals must modify their behavior based on updated expected outcomes in a changing environment. Prelimbic cortex (PrL) neural encoding during learning predicts and is necessary for appropriately altering behavior based on new expected outcome value following devaluation. We aimed to determine how PrL neural activity encodes reward predictive cues after the expected outcome value of those cues is decreased following conditioned taste aversion. In one post-devaluation session, rats were tested under extinction to determine their ability alter their behavior to the expected outcome values (i.e., extinction test). In a second post-devaluation session, rats were tested with the newly devalued outcome delivered so that the rats experienced the updated outcome value within the session (i.e., re-exposure test). We found that PrL neural encoding to the cue associated with the devalued reward predicted the ability of rats to suppress behavior in the extinction test session, but not in the re-exposure test session. While all rats were able to successfully devalue the outcome during conditioned taste aversion, a subset of rats continued to consume the devalued outcome in the re-exposure test session. We found differential patterns of PrL neural encoding in the population of rats that did not avoid the devalued outcome during the re-exposure test compared to the rats that successfully avoided the devalued outcome. Our findings suggest that PrL neural encoding dynamically tracks expected outcome values, and differential neural encoding in the PrL to reward predictive cues following expected outcome value changes may contribute to distinct behavioral phenotypes.

## 1.1 Introduction

The ability to alter to behavior in response to changes in consequences is necessary for navigating an ever-changing environment. Many neuropsychiatric disorders are characterized by a continuation of maladaptive behavior in spite of negative consequences [1, 2]. Thus, characterizing the underlying processes that modulate the ability to change or stop behavior in response to updated expected outcomes is critical for understanding the neurobiological alterations in neuropsychiatric disorders. The ability to use new updated expected outcome values to direct behavior can be measured using outcome devaluation tasks. There are several necessary stages to perform this task: 1) forming an association between a cue and an outcome, 2) registering the devalued outcome, and 3) integrating the cue-outcome association with the decreased outcome value to guide behavior. During this stage, animals typically do not experience the cue together with the now devalued outcome because testing is performed under extinction in rodents. As such they must rely on the internal representations of the originally established cue-outcome association and the subsequent devaluation of the outcome to successfully alter behavior [3, 4], a complex feat that requires the functional integration of frontal neural networks. Finally, there are individual differences in both animals and humans in how they encode these distinct computational processes, and these differences may underlie differing susceptibility across individuals to neuropsychiatric disorders [5-7] suggesting the broader utility of the outcome devaluation task in animal models.

The rat frontal cortex has been implicated in modulating behavior based on newly computed outcome values. The prelimbic cortex (PrL) neurons show increased responsiveness to cues following cue-outcome associative learning [8-10]. In addition, PrL is necessary during learning (i.e., forming associations) to guide subsequent goal-directed behavior [11-14] following outcome devaluation. However, the PrL is not necessary when animals must update outcome value during the post-outcome devaluation test session. This suggests that PrL is not critically involved in integrating cue-outcome associations with the devaluation of the outcome to ultimate modify behavior. This seems incongruent with other tasks that involve modified expected outcome values during decision-making. For example, PrL plays a key role in integrating information about updating value representations for optimal choices in value-based decision-making tasks [15-18]. PrL encoding is linked to the evaluation of negative outcomes to appropriately adjust responding for optimal risky decision-making [18] and also tracks preferred reward value during delay based decision-making [17].

One distinction between the outcome devaluation task and value-based decision-making tasks is that in valued-based decision-making tasks, rats continue to earn the expected outcome throughout the session. As such, rats are not required to use internal representations of those values to guide their behavior, i.e., the expected outcome value shifts within the session, and the rats can reform associations within a behavioral session. In contrast, in outcome devaluation tasks, rats never experience the new updated outcome value with the cue, i.e. they do not receive real-time feedback to guide behavior which would allow them to learn to avoid the cue if they were unable to integrate previously learned information [3, 4]. Thus, the seeming discrepancy between the role of the PrL during the integration of updated expected outcome value during outcome devaluation tasks and value-based decision making may be related to whether animals have the opportunity to learn and update their representations about the value in real-time. PrL neural encoding dynamically updates outcome value during set-shifting in real-time where the outcomes are delivered through the session [19, 20], supporting its potential role in flexibility with feedback. We previously have shown that PrL neural encoding throughout learning predicts how well rats are able to suppress their behavior post-devaluation [14]. Here, we aimed to determine how the PrL encodes information related to reward predictive cues after the expected outcome value of those cues has been decreased, i.e., when the animals alter their behavior in real-time to adjust to those expected outcome values compared to when the animals experience the updated outcome value within the session. In addition, we aimed to determine if re-introducing the outcome post-devaluation would affect behavioral performance and/or PrL neural encoding in this task.

## 2. Materials and Methods

### 2.1. Subjects

Long-Evans male rats (n=20), aged 90-120 days (250-350 grams) at the beginning of the study were used. All subjects were housed individually and maintained on a standard 12:12 hour light-dark cycle (lights off at 7:00 am). During behavioral training and testing, rats were maintained at no less than 90% of their preoperative body weight by regulating food access to 20-25 g of standard rat chow per day (*ad libitum* water). All animal procedures were approved by the University of North Carolina at Chapel Hill and the Rowan University School of Osteopathic Medicine Institutional Animal Care and Use Committees (IACUC). We have previously reported *in vivo* electrophysiology data during Pavlovian conditioning (not presented here) from a subset of the rats (n=16) in this study [14]. We have not previously reported any of the *in vivo* electrophysiological data from our post-devaluation test days represented in this study (see below). The timeline of this experiment is shown in Figure 1A.

**Figure 1.**
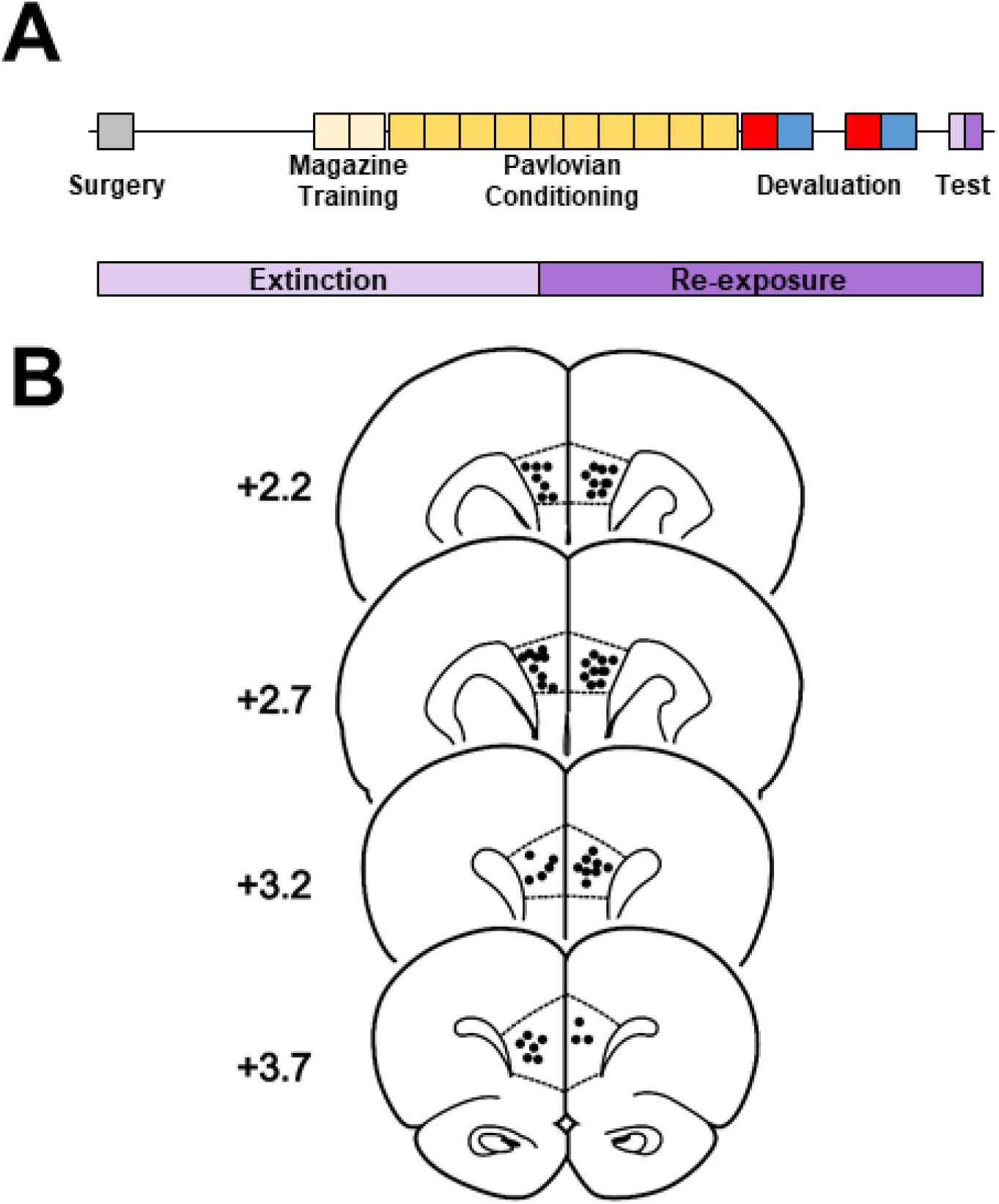
A) Time of experiments including surgery, magazine training (2 days), Pavlovian conditioning (10 days), devaluation (sugar pellets/LiCl, red square and food pellets/saline, blue square), and test day (bottom, representation extinction test followed by re-exposure test). B) Histological verification of recording array wires in the PrL. Filled circles indicate electrode locations in the PrL. The numbers represent distance in millimeters from bregma.

### 2.2. Electrophysiological implant surgery

Rats were anesthetized with a ketamine hydrochloride (100 mg/kg) and xylazine hydrochloride (10 mg/kg) cocktail (i.p.).Rats surgically implanted with microwire arrays (8 microwires per implant) in the PrL (AP +2.5, ML, ±0.6 mm, DV -5.0 from skull [14, 17, 21]).

### 2.3. Behavioral Task

Rats were trained on a conditioned outcome devaluation task consisting of three phases: Pavlovian conditioning, devaluation and two post-devaluation tests (Figure 1A). First, rats underwent two days of magazine training where 20 each of sugar and food pellets were delivered. During Pavlovian conditioning, one CS^+^ consisted of the illumination of two cue lights located to the left and right sides of the food receptacle (i.e., solid CS^+^). The other CS^+^ consistent of the flashing illumination (5 Hz) of the two cue lights located to the left and right side of the food receptacle (i.e., flashing CS^+^). The food paired with the two CS^+^ was counterbalanced such that a subset of rats received the food pellets paired with the solid CS^+^ (and the sugar pellets paired with the flashing CS^+^) and the other half received the reverse. Importantly, rats were also exposed to two distinct CS^-^ consisting of either a solid or flashing cue light located above the food receptacle (i.e., solid CS^-^ and flashing CS^-^). The CS^+^ paired with food (regardless of solid or flashing) is designated as CS^+1^ and the CS^+^ paired with sugar is designated as CS^+2^. Each rat received 10 presentations of each stimuli and the order of the presentation were pseudorandomized. The inter-trial interval for the stimuli was pseudorandom and variable (75, 90, 105, or 120 seconds). There were three different sessions with different stimuli presentation order, however, the same order was used for Day 1, Day 10, and the post-devaluation tests.

During the devaluation phase, rats were habituated for two days in a standard empty rat cage which would be used for the subsequent devaluation procedure (30 mins/day: on the last two days of Pavlovian conditioning,). After habituation, rats began the devaluation procedure. Here, rats were allowed 30 minutes to eat one of the rewards (food or sugar) *ab libitum* in the empty cage. Following consumption of the sucrose pellets, rats received an injection of LiCl (0.3 M, 7.5 ml/kg, i.p. [14, 22] while following consumption of the food pellets rats received an injection of saline (i.p.). At least 48 hours later, the same two-day procedure was repeated. At least another 48 hours after the completion of devaluation, rat were run through two post-devaluation test sessions. For the first test session, rats were given the same paradigm in Pavlovian conditioning (10 presentations of each stimuli, CS^+1^, CS^+2^, CS^-1^, CS^-2^), however, no reinforcer was delivered during testing (extinction test) during which rats must use expected outcome to guide behavior. Following this extinction test session, rats underwent another Pavlovian session accompanied by the reintroduction of both reinforcers (i.e., now nondevalued and devalued; re-exposure test) in which rats can use the actual outcome value to guide behavior. In this study we exclusively devalued the sucrose pellets as the food pellets were similar to what the rats received in their home cages and we wanted to avoid interfering with their daily food intake. Critically, we have previously shown that the devaluation of multiple reinforcers can be transferred to the reward predictive cues using similar methods [23-26].

### 2.4. Electrophysiological Recordings

Online isolation and discrimination of single unit activity was accomplished using a commercially available neurophysiological system (OmniPlex system; Plexon). Continuous recordings from electrodes were virtually referenced (PlexControl, Plexon) and fed into a Pentium computer. Continuous signals were high-pass filtered (300Hz) to identify individual spike events. Discrimination of individual waveforms began by setting a threshold level (3.5σ) for each wire. Individual waveforms corresponding to a single cell were discriminated using template analysis procedures and time–voltage boxes provided by the neurophysiological software system. Cell recognition and sorting was finalized after the experiment using the Offline Sorter program (Plexon, Inc). This allowed neuronal data to be further assessed based on the principle component analysis of the waveforms, cell firing characteristics such as autocorrelograms and interspike interval distribution to ensure that putative cells showed biologically appropriate firing refractory periods, and cross-correlograms to ensure that multiple cells recorded on the same wires fired independently of each other. Waveform and spontaneous firing rates were examined to identify putative glutamatergic pyramidal neurons in the PrL in the analysis [10]. Finally, an additional computer processed operant chamber input and output (Med Associates, Inc) and sent digital outputs corresponding to each event into system to be time stamped along with the neural data.

### 2.5. Data Analysis

#### 2.5.1. Behavior

We assessed the degree of learning across Pavlovian conditioning using a repeated measures two-way ANOVA with day (Day 1 vs Day 10) and cue presentation (CS+1, CS+2, CS-1, CS-2) as factors. To confirm successful devaluation, we analyzed the amount of food consumed during the LiCl procedure using repeated measures (RM) two-way ANOVA with pairing (first pairing vs second pairing) and outcome (to be nondevalued: food vs to be devalued: sugar) as factors. To assess the ability of rats to shift behavior during the post-devaluation tests, we compared percent of time spent in the food cup during each CS+ (devalued, nondevalued) on the post devaluation test (extinction test and re-exposure test sessions) using a pair student’s t-test comparing devaluation status (nondevalued vs devalued). We also calculated a devaluation index (DI) using the following formula: (CS^+^ predicting nondevalued (ND) outcome - CS^+^ predicting devalued (D) outcome)/ (CS^+^ predicting nondevalued (ND) outcome + CS^+^ predicting devalued (D) outcome)/ or (ND-D)/(ND+D) as we have previously done[14, 23, 25, 26]. Based on this formula, the greater the rat suppressed approaching the cup in the devalued condition (but continued approaching the cup in the nondevalued condition), the closer the value would be to 1. The ability to suppress approach behavior towards the CS+ that predicts the devalued reinforcer (DI values > 0) represented flexible behavior. We used these values to correlate the strength of the devaluation effect (DI, - 1 to 1) with the % of phasic responsive in PrL neurons in individual rats during both post-devaluation tests (extinction and re-exposure). Finally, based on the finding that two distinct population of rats emerged with regard to the number of pellets rats consumed during the post-devaluation exposure test, we analyzed the devaluation data and post-devaluation test days using RM two-way ANOVA with devaluation status as a within subject factor (nondevalued vs devalued) and a between subject factor of avoidance type (Avoider vs Non-Avoider). The conditioned taste aversion consumption data was analyzed using a RM-three way ANOVA using within subject factors, outcome (to be nondevalued: food vs to be devalued: sugar), pairing (1^st^ vs 2^nd^), and between subject factor of avoidance type (Avoider vs Non-Avoider).

#### 2.5.2 Electrophysiology

Changes in neuronal firing patterns relative to task events were analyzed by constructing peri-event histograms (PEHs) surrounding each cue presentation using commercially available software (Neuroexplorer for Windows version 4.034; Plexon, Inc.). PEHs (200-ms bins; 20 s total) were constructed on the post-devaluation test day to the cues that predicted the nondevalued outcome and the cues that predicted the Devalued outcome. The activity of each cell was examined relative to cue (0–10 s following cue presentation) for all trials. Individual units were categorized as showing either a decrease (inhibition) or an increase (excitation) in firing rate compared to baseline (i.e., termed ‘phasic’ activity) or no difference in activity from baseline (termed ‘nonphasic’). Specifically, cells were classified as phasic if during cue presentation the firing rate was greater than (excitation) or less than (inhibition) the 99.9% confidence interval projected from the baseline period for at least one 200-ms time bin [14, 21, 23, 27]. This confidence interval was selected such that only robust responses were categorized as excitatory or inhibitory [14]. Some neurons in this analysis exhibited low baseline firing rates and the 99.9% confidence interval included zero. When this was the case, inhibitions were assigned if the number of consecutive 0 spikes/0 bins in the event epoch was at least double the number of consecutive 0 spikes/s time bins during the baseline period [14, 21, 23, 27]. Units that exhibited both excitations and inhibitions within the same epoch were classified by the response that was most proximal to the event. The percentage of phasic cells that responded to the cue that predicted the nondevalued vs the cue that predicted the devalued outcome in the PrL in each individual animal was calculated for both post-devaluation test sessions (extinction and re-exposure test). We analyzed the percentage of phasic neurons in nondevalued vs devalued conditions using a paired t-test. In addition, each individual animal’s percentage of phasic neurons to the nondevalued and devalued cues were correlated with the animals’ subsequent test day performance as measured by the Devaluation Index (see above). Only animals with at least three cells recorded on the test day were included in this analysis (n=16).

### 2.6. Histology

Upon completion of the experiment, rats were deeply anesthetized with an intraperitoneal injection of a ketamine and xylazine mixture (100 and 10 mg/kg, respectively). For *in vivo* electrophysiology studies, a 13.5-μA current was passed through each microwire electrode for 5 s to mark the placement of electrode tips. Transcardial perfusions were then performed using physiological saline and 3% potassium ferricyanide in 10% formalin, and brains were removed. After post-fixing and freezing, 40-μm coronal brain sections were mounted. The addition of potassium ferricyanide allowed for a blue reaction corresponding to the location of the electrode tip which was viewed under a 1X microscope lens. Placement of an electrode tip within PrL was determined by examining the relative position of observable reaction product to visual landmarks and anatomical organization of the PrL represented in a stereotaxic atlas (Figure 1B).

## 3. Results

### 3.1. Prelimbic neural encoding in real-time predicts the ability to suppress behavior following a decrease in expected outcome value

We recorded from 106 neurons in the PrL on the two post-devaluation test sessions (extinction and re-exposure) with the breakdown of our wire placements shown in Figure 1.

We found that PrL neural encoding dynamically tracks expected outcome value and predicts the ability of rats to suppress approach behavior post-devaluation. After 10 days of Pavlovian conditioning, rats spend more time in the food cup to both CS+1 and CS+2 compared to CS-1 and CS-2 (Figure 2A) as revealed by a two-way Repeated Measures (RM) Analysis of Variance (ANOVA) (Day: F_(1,18)_=81.7, CS: F_(3,54)_=46.2, Day x CS interaction: F_(3,54)_=45.1, Tukey’s post-hoc). Rats successfully devalue the sugar pellets after one pairing with LiCl pairing while continuing to eat the food pellets that were paired with saline (Figure 2B) as revealed by a two-way RM-ANOVA (Pairing: F_(1,19)=_32.1, Outcome: F_(1,19)_=17.2, Pairing x Outcome interaction, F_(1,19)_ =42.5, Tukey’s post-hoc). As shown in Figure 2C, rats go into the foodcup less during the presentation of cue paired with sugar (devalued) compared to the cue paired with food (nondevalued) during the post-devaluation extinction test (t_(19)_=3.5, p<0.05). This suggests that rats were able to successfully update the information about the devalued outcome to suppress their behavior towards the cue that predicted the devalued outcome, while continuing to respond to the cue predicting the nondevalued outcome. Distinct PrL neurons show “phasic” responsiveness to the cues that were classified as either excited or inhibited relative to presentation of the cue during the post-devaluation test (Figure 2D-E). We found that the percentage of phasic neurons (excited or inhibited response to the cue) in individual rats is significantly less to the cue paired with sugar (devalued) compared to the cue paired with food (nondevalued) during the post-devaluation extinction test (t_(15)_=2.8, p<0.05) as shown in Figure 2F. In addition, we found a significant negative correlation between the percentage of phasic responsive PrL neurons to the devalued cue and behavioral performance measured as a devaluation index (Figure 2G), but no correlation with the nondevalued cue and performance. Interestingly, the percentage of PrL neurons that showed an excited profile (Figure 2H) to the devalued cue negatively correlated with behavioral performance, while we found no correlation between the PrL neurons with inhibited profile and behavioral performance (Figure 2I). We also found no correlation of excited or inhibited neurons to the nondevalued cue and behavioral performance (Figure 2H-I). Together these data suggest that the more PrL neurons are engaged to the cue that predicts the expected devalued outcome (i.e., greater population of phasic responsive neurons, particularly “excited”) the worse the rat’s ability to suppress behavior towards the devalued cue (i.e., during the post-devaluation extinction test).

**Figure 2:**
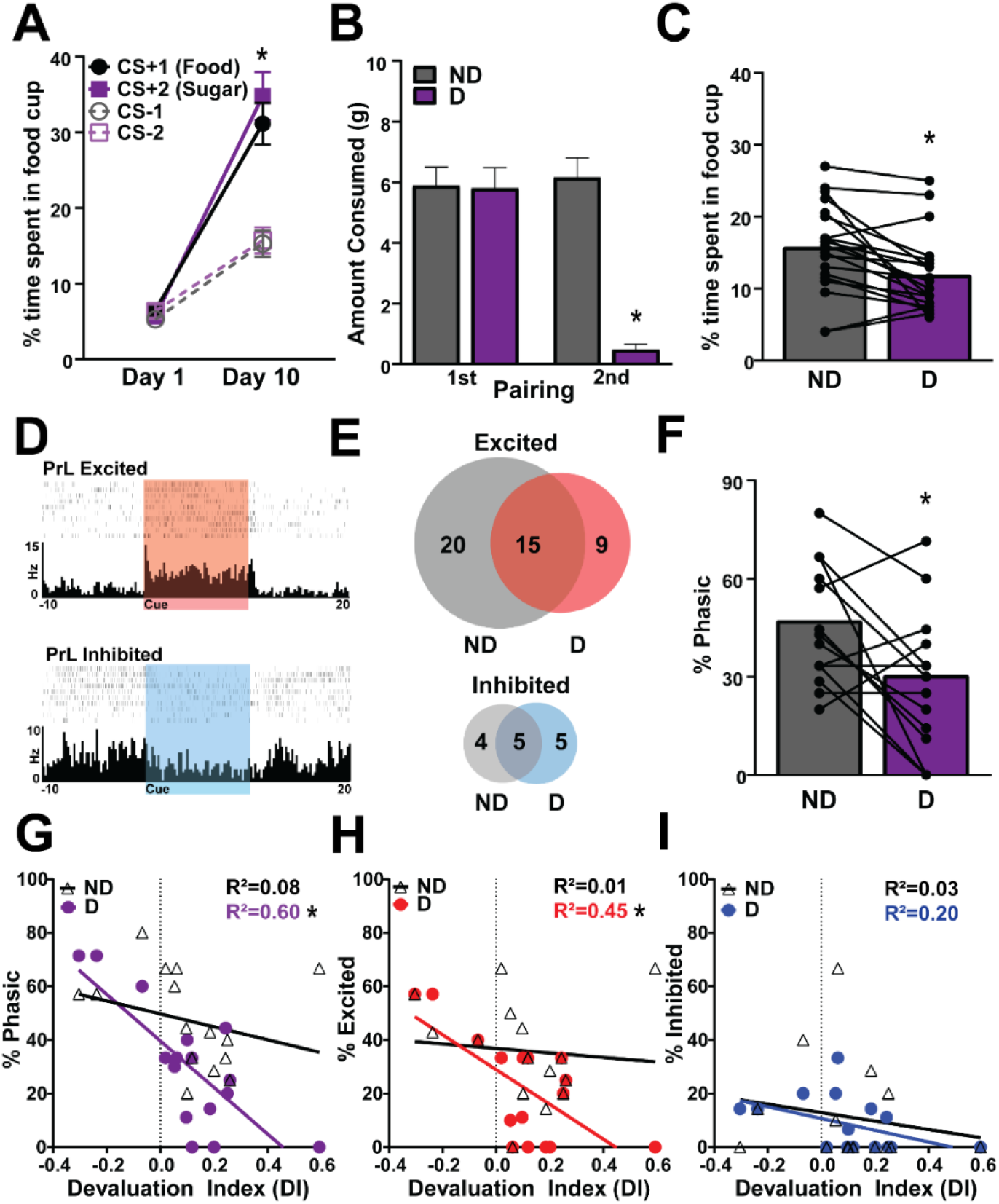
PrL neural encoding dynamically tracks expected outcome value and predicts the ability of rats to suppress approach behavior during the post-devaluation extinction test. A) After 10 days of Pavlovian conditioning, rats spend more time in the food cup to both CS+1 and CS+2 compared to CS-1 and CS-2. B) After one sugar/LiCl pairing rats decrease the amount of sugar they eat showing successful sugar devaluation for second pairing. Lines represent individual rats. C, Rats respond less to the CS^+^ paired with sugar (devalued) compared to the CS^+^ paired with the nondevalued outcome post-devaluation extinction test, lines represent individual rats. D, Perievent histograms showing examples of individual “phasic” PrL neurons that are classified as either excited (top, red) or inhibited (bottom, blue) relative to presentation of the cue during the post-devaluation test. E) Venn diagrams of the population response of PrL neurons to the nondevalued cue, devalued cue, or the overlap for PrL neurons classified as excited (top, red) or inhibited (bottom, blue) during the post-devaluation extinction test, 48 neurons were classified as nonphasic. F) The percentage of phasic neurons (excited or inhibited response to the cue) in individual rats was significantly less to the cue paired with sugar (Devalued, D) compared to the cue paired with nondevalued outcome post-devaluation, lines represent individual rats. G) Percentage of PrL neurons that exhibited phasic responsiveness during cue presentation to either the nondevalued cue (white triangles) or devalued cue (purple circles) plotted against each rat’s devaluation index. There was a significant negative correlation between percentage PrL phasic neurons to the devalued cue, but not the nondevalued cue, and devaluation index. Percentage of PrL neurons that exhibited an excited response during cue presentation to either the nondevalued cue (white triangles) or devalued cue (red circles) plotted against each rat’s devaluation index (extinction test). There was a significant negative correlation between percentage excited PrL neurons to the devalued cue, not the nondevalued cue, and devaluation index (extinction test). I) Percentage of PrL neurons that exhibited an inhibited response during cue presentation to either the nondevalued cue (white triangles) or devalued cue (blue circles) plotted against each rat’s devaluation index (extinction test). There was no correlation in the percentage of inhibited PrL neurons to the devalued or nondevalued cue and devaluation index (extinction test). CS= conditioned stimulus, PrL= prelimbic cortex, ND=nondevalued, D=devalued. Mean +/-SEM, * p<0.05

### 3.2. Prelimbic neural encoding does not dynamically change to cues that predict the devalued outcome

During the subsequent post-devaluation re-exposure test (Figure 1B), we found that rats spend less time in the food cup during the cue paired with sugar (devalued) compared to the cue paired with food (nondevalued; Figure 3A, t_(19)_=2.4, p<0.05). Rats ate fewer sugar pellets (devalued) compared to food pellets (nondevalued) during the post-devaluation re-exposure test following outcome delivery (Figure 3B, t_(19)_=4.03, p<0.05). In addition to classifying neurons as phasic to the cue (as in Figure 2D), we also classified individual “phasic” PrL neurons as either excited or inhibited to the outcome delivery (Figure 3C) during the re-exposure test session. We found similar PrL population responses to the nondevalued and/or devalued to the cues or outcomes during the post-devaluation re-exposure test (Figure 3D). We also found that there were no differences in the percentage of phasic neurons (excited or inhibited response) to the nondevalued and devalued cue or the nondevalued and devalued outcome (Figure 3E) as revealed by a two-way RM ANOVA (Cue: t_(15)_=0.25, p>0.1, Outcome: t_(15)_=0.38 p>0.1). In contrast to our PrL neural encoding during the extinction test, we found no correlation between the percentages of PrL neurons that exhibited phasic responsiveness during cue presentation or the outcome relative to each rat’s devaluation index during the post-devaluation re-exposure test (Figure 3E). Thus, PrL neural activity during the post-devaluation extinction test predicted performance during this test session, while PrL neural activity during the re-exposure test had no predictive value to behavioral performance during the test session. Based on this finding, we aimed to determine if there were differences based on whether or not the rats ate the outcomes during the re-exposure test.

**Figure 3:**
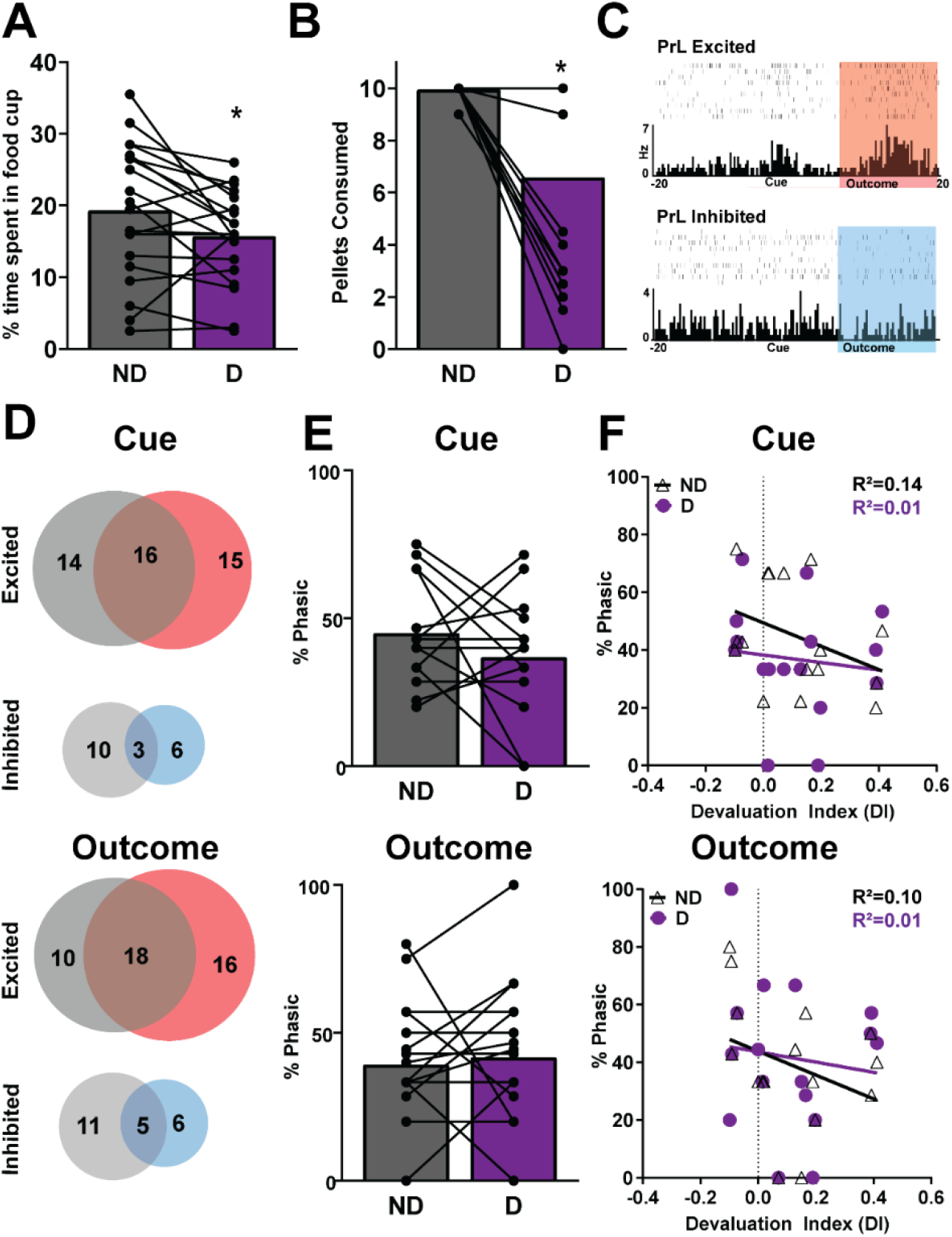
PrL neural encoding does not track neural encoding based on actual outcomes to their predictive cues or outcomes during a post-devaluation re-exposure test. A) Rats respond less to the CS^+^ paired with sugar (Devalued, D) compared to the CS^+^ paired with food (NonDevalued, ND) during the post-devaluation re-exposure test, lines represent individual rats. B) Rats eat fewer sugar pellets (Devalued, D) compared to food pellets (NonDevalued, ND) during the post-devaluation re-exposure test following outcome delivery, lines represent individual rats. C, Perievent histograms showing examples of individual “phasic” PrL neurons that are classified as either excited (top, red) or inhibited (bottom, blue) relative to outcome delivery during the post-devaluation re-exposure test. D) Venn diagrams of the population response of PrL neurons to the cues (top) or outcome (bottom) represent to the nondevalued, devalued, or the overlap for PrL neurons classified as excited (top, red) or inhibited (bottom, blue) during the post-devaluation exposure test, 45 neurons were classified as nonphasic to the cue and 38 neurons were classified as nonphasic to the outcome. E) The percentage of phasic neurons (excited or inhibited response) to the cue (top) or the outcome (bottom) in individual rats is no different in the Devalued (D) condition compared to NonDevalued (ND) during the post-devaluation re-exposure test lines represent individual rats. F) Percentage of PrL neurons that exhibited phasic responsiveness during cue presentation (top) or the outcome (bottoms) in either the nondevalued condition (white triangles) or devalued condition (purple circles) plotted against each rat’s devaluation index during re-exposure test. There were significant no correlations in PrL encoding to either the nondevalued cue or devalued cue (top) and devaluation index (re-exposure test) and no correlations to either the nondevalued outcome or devalued outcome (bottom) and devaluation index (re-exposure test). CS= conditioned stimulus, PrL= prelimbic cortex, ND=nondevalued, D=devalued

### 3.3. Distinct phenotypes emerge with differential behavioral patterns and PrL neural activity cue that predict expected outcome value

We observed that during the post-devaluation re-exposure test, a subset of rats continued to consume the devalued outcome in spite of it being previously paired with LiCl. We found that 11 rats continued to eat the sugar pellets (consumed: 9.8 +/-0.8) and that 9 rats avoided the sugar (consumed: 2.6 +/-1.4) as such the mean for each group of rats was greater than two standard deviations from each other representing a bimodal distribution as shown in Figure 4A. We re-analyzed our behavioral and in *vivo* electrophysiological data separating the groups based on these two distinct behavioral phenotypes. Rats that successfully avoided the sugar pellets were classified as “Avoiders” and rats that continued to consume the sugar pellets were classified as “Non-Avoiders”. Critically, rats that were classified as Avoiders and Non-Avoiders were both able to successfully devalue the sugar after pairing with LiCl, while continuing to consume the alternative, nondevalued, outcome (Figure 4B) as shown by a Three-way RM ANOVA (Avoider vs Non-Avoider, F_(1,14)_=0.10, Pairing 1 vs Pairing 2, F_(1,14)_=32.6, nondevalued vs devalued F_(1,14)_=19.6, Pairing x Devaluation Status interaction F_(1,14)_=26.0, Bonferroni post-hoc revealed pairing 2 for the devalued outcome different from pairing 1 for both outcomes and pairing 2 for nondevalued outcome in both Avoiders and Non-Avoiders).

**Figure 4.**
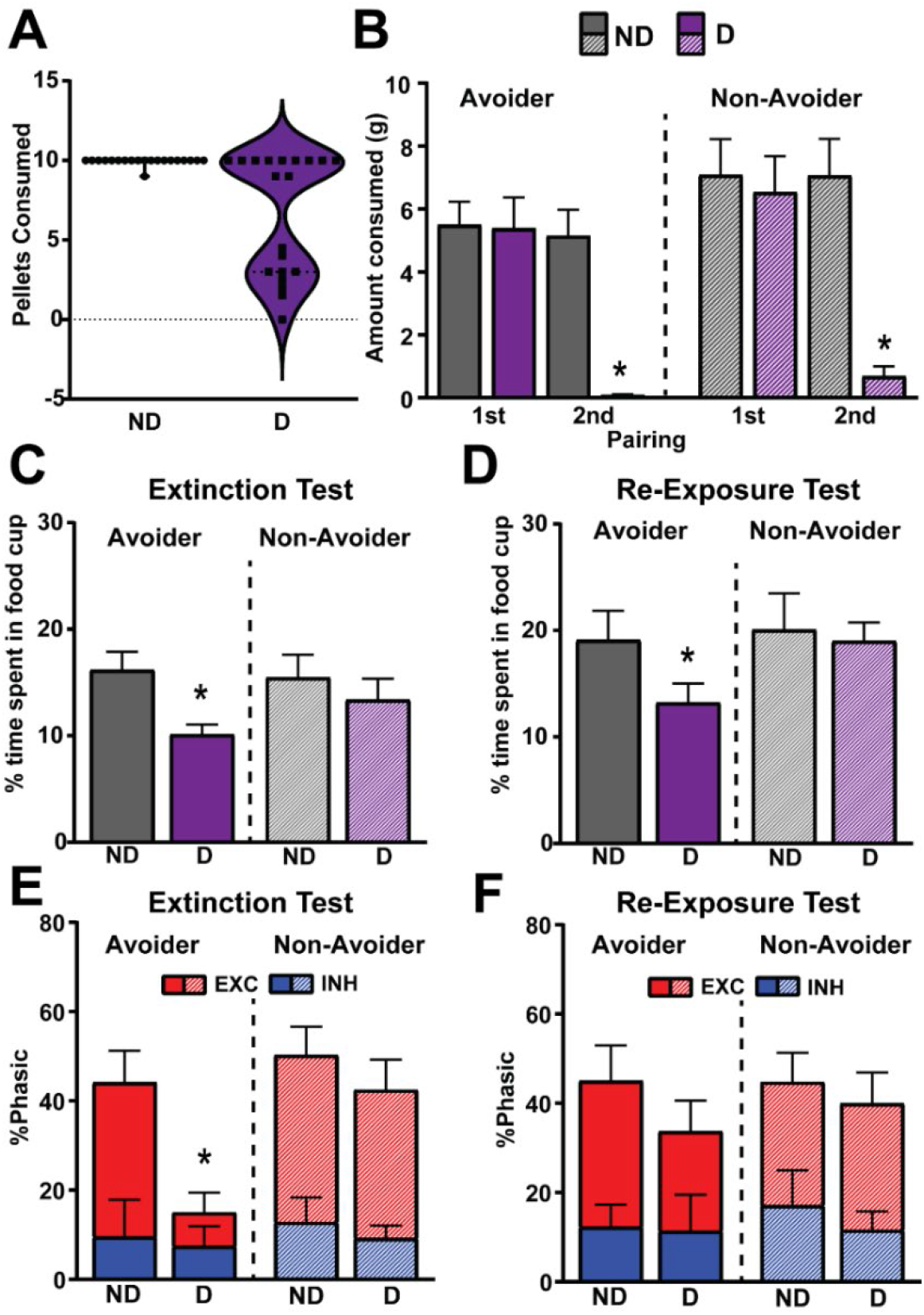
A) Differential behavioral responses and PrL encoding in rats that successfully avoided (Avoider) the devalued outcome and rats that failed to avoid the devalued outcome (Non-Avoider) in the post-devaluation re-exposure test. A) Two distinct populations of rats are represented in the violin plot in relation to how much they eat of the devalued outcome. B) Both groups of rats (Avoiders and Non-Avoiders) show significant devaluation effect to the devalued outcome for the 2^nd^ pairing but not the nondevalued outcome (* p<0.05 different from nondevalued 2nd pairing and devalued 1^st^ pairing) following conditioned taste aversion and there are no differences between these two groups. C-D) Rats that were classified “Avoiders” spent less time in the food cup to the cue the predicted the nondevalued outcome (ND) compared to the Devalued (D) outcome, while rats that were classified as “Non-Avoiders” spent equal time in both cues in the post-devaluation (C) extinction test (ND vs D, *p <0.05). E) Rats that were classified “Avoiders” showed significantly less phasic responsiveness, specifically excited responses, to the cue that predicted the NonDevalued outcome (ND) compared to the cue that predicted that the Devalued (D) outcome, while rats that were classified as “Non-Avoiders” showed equal phasic responsiveness to both cues during the post-devaluation extinction test (ND vs D, *p <0.05) F) Neither group of rats (Avoiders or Non-Avoiders) showed any differences in the percentage of phasic responsiveness to the cues during post-devaluation re-exposure test. PrL= prelimbic cortex, ND=nondevalued, D=devalued.

In the post-devaluation extinction test, rats that were classified Avoiders spent less time in the food cup to the cue the predicted the nondevalued outcome compared to the devalued outcome, while rats that were classified as Non-Avoiders did not (Figure 4C) as revealed by a Two-way ANOVA (Avoider vs Non-Avoider: F_(1,18)_=0.03, nondevalued vs devalued: F_(1,18)_=15.1, group by devaluation status interaction, F_(1,18)_=3.6, Bonferroni post-hoc revealed a significant different between time spent in the cup during the nondevalued cue and devalued cue only in Avoiders). Similarly, in the post-devaluation re-exposure test, Avoiders, but not Non-Avoiders, spent less time in the food cup to the cue the predicted the nondevalued outcome compared to the devalued outcome (Figure 4D) as revealed by a Two-way RM ANOVA (Avoider vs Non-Avoider: F_(1,18)_=1.1, nondevalued vs devalued: F_(1,18)_=5.0, group by devaluation status interaction, F_(1,18)_=2.4, Bonferroni post-hoc revealed a significant difference between time spent in the cup during the nondevalued cue and devalued cue only in the Avoider group). Further, rats that were classified “Avoiders” showed significantly less phasic responsiveness to the cue that predicted the devalued outcome compared to the cue that predicted that the nondevalued outcome, while rats that were classified as Non-Avoiders did not show this difference during the post-devaluation extinction test (Figure 4E) as shown by a Two-way RM ANOVA (Avoider vs Non-Avoider: F_(1,14)_=4.8, nondevalued vs devalued: F_(1,14)_=6.5, group by devaluation status interaction, F_(1,14)_=2.3, Bonferroni post-hoc revealed a significant difference in PrL phasic responsiveness to the nondevalued cue and devalued cue only in the Avoider group). Furthermore the diminished phasic responsiveness to the devalued cue in Avoiders is primarily due to a decrease in the neurons that are classified as excitations (Figure 4E). In contrast, neither group of rats (Avoiders or Non-Avoiders) showed any differences in the percentage of phasic responsiveness to the cues during post-devaluation re-exposure test as shown by a Two-way RM ANOVA (Avoider vs Non-Avoider: F_(1,14)_=0.2, nondevalued vs devalued: F_(1,14)_=1.3, group by devaluation status interaction, F_(1,14)_=0.21).

## 4. Discussion

Our findings suggest that PrL dynamically updates and integrates the expected outcome value to the predictive cue during the post-devaluation extinction test session, but does not show differential neural encoding to reward predictive cues that predict the new updated actual outcome value. This is consistent with PrL encoding shifts in expected value in value-based decision making tasks [17, 19] even though the PrL is not necessary during the test sessions following outcome devaluation while rats shift their behavior[12]. This could be due to differing task structure (e.g., Pavlovian vs Instrumental) or it could result from other brain regions compensating for the loss of PrL function during the task (e.g., orbitofrontal cortex and/or basolateral amygdala [28-33]). Interestingly, the PrL neural encoding in this task is distinct from what we have previously observed in the NAc core [23]. NAc core, a prime target of PrL glutamatergic output [34], neural activity does not change with regard to decreased expected outcome value when rats must use the internal representations to guide behavior as the decreased outcome value and the reward are never paired [23], even though NAc neurons encode relative outcome values and subsequent behavioral responses when rats have the opportunity to updated the expected outcome value in real-time[27, 35, 36]. One possibility is that the NAc is more involved in experiential updating of actual outcome value. This would be consistent with the idea that the NAc core specifically is involved in learning (or re-learning) cue-outcome associations [37-39].

Importantly, this dynamic encoding of PrL is specific to the subset of rats that avoided the devalued outcome in re-exposure test (Avoiders). The rats that continued to consume the devalued outcome in post-devaluation re-exposure test (Non-Avoiders) continued to show elevated behavioral and PrL neural responding to both the nondevalued and devalued condition in both post-devaluation test sessions. These findings suggest that Avoider group was better at using the updated expected outcome value to guide behavior during the preceding post-devaluation extinction test. This was likely not due to a decrease in behavioral responding since rats that were classified as Avoiders also spent less time in the food cup to the cue that predicted the devalued outcome in the re-exposure test, even though the PrL showed elevated phasic responsiveness during this test session.

Critically, the classification of the “Avoider” vs “Non-Avoider” phenotypes was based on the consumption of the nondevalued vs devalued outcome during post-devaluation re-exposure test when the rats were in the operant chamber undergoing conditioning, i.e., cues predicted the outcome delivery. This was not an operational measure as to whether they avoided or did not avoid the sugar during devaluation procedure itself (i.e., all rats avoid the devalued outcome outside of the conditioning task). This suggests that this Non-Avoider subset of rats were unable to stop engaging in the pre-learned response to the CS+ to continue eating the outcome (cue light on, go to food cup, eat outcome) in spite of the outcome being devalued, even though these rats showed successful devaluation. We previously reported a similar phenomenon in monkeys following orbitofrontal cortex inactivation in which they continued to eat a devalued outcome after the monkey removed the object paired with that outcome[24], in spite of the monkey successfully avoiding the devalued outcome when not required to initiate the motor sequence response to the conditioned stimulus (remove object to retrieve outcome). This is also consistent with our previous observations that a subset of rats are insensitive to outcome devaluation without any pharmacological or neural perturbations, and our current findings further characterize these phenotypes [14, 23].

While the PrL neurons show less engagement to the devalued cue in the post-devaluation extinction test (in Avoiders), the PrL is re-engaged during the post-devaluation re-exposure test.

One possibility is that the PrL becomes reactivated as rats are re-learning information about the cue-outcome associations particularly given the role of PrL in mediating attention [19, 20, 40]. This would be consistent with previous work showing some PrL neurons are activated by changing rule contingencies[19]. PrL neural activity predicts behavior more accurately later within a set shifting task, suggesting that the activity likely is more refined as the updating behavior solidifies [20]. In our re-exposure test session, the rats are likely starting to reform their associations, as such PrL activity is elevated in both the nondevalued and devalued conditions. Our findings that there is no predictive activity of the PrL activity during the outcome delivery itself is consistent with previous work showing that neural encoding of the PrL is outcome predictive but is not responsive to outcomes[20]. If the PrL is not responsive to outcomes in general, but it is responsive to attention and shifting behavior, then perhaps the PrL is re-engaged in the re-exposure test because the rats are attending to both the cues and their updated outcomes. In addition, the PrL is involved in fear learning [41-43], so the PrL may be re-engaged in this session, particularly in the Avoider phenotype, as they are re-establishing that the cue predicts a negative outcome. Thus, while we observe a similar neural phenotypes in the Avoiders and Non-Avoiders, they may represent distinct processing within the PrL. For example, perhaps in the Non-Avoiders the engaged PrL is representative of these rats continuing to respond to both the nondevalued and devalued. While, in the Avoider group, the rats are showing elevated engagement due to paying closer attention to the cues and/or forming the new association with the cue and the new negative outcome.

This very clear bimodal phenotype could have further implications related to individual susceptibility to other perturbations that affect the frontal cortex such as in Substance Use Disorders[44] or anxiety disorders [2]. For example, a history of drug use alters learning and valued based decision-making [45-47] but also distinct deficits in learning and decision-making are linked to increased susceptibility to drug seeking or escalation of drug use[7, 48]. Critically, a history of psychostimulants[21, 23, 49, 50], opioids[51], ethanol[52], or stress [53-56] alters frontal neural activity and disrupt learning and decision-making. A rodent model of obsessive-compulsive disorder shows elevated PrL neural activity and impairments in reversal learning [57]. Our current findings taken with animal models of neuropsychiatric disease suggest that PrL neural activity may need to be optimally dynamic for flexible behavior as impaired behavior has also been reported when PrL neural activity is dampened [14, 51] or heightened [57]. Finally, it would be interesting if rats with one of the distinct phenotypes are more susceptible to drug seeking or more vulnerable to stressors given that rats that show deficits in behavioral flexibility measured by probabilistic reversal learning are more likely to escalate drug seeking [6]. As such future studies will aim to determine if these distinct phenotypes and underlying PrL encoding are linked to subsequent maladaptive behavior (e.g., are animals classified as the Non-Avoider phenotype more likely to engage in escalating drug seeking or risky decision-making?).

## 5. Conclusions

## Acknowledgements

This work was funded by the National Institute on Drug Abuse (R00DA042934 to EAW).

## Notes

### Competing Interest Statement

The authors have declared no competing interest.

